# Commensal enteric virome regulates intestinal carbohydrate sensing via Th17 cells

**DOI:** 10.1101/2024.10.15.618587

**Authors:** Fengqing Lin, Lianglan Li, Yuanpeng He, Shujie Xu, Qiufen Mo, Wangsen Cao, Aikun Fu, Weiqin Li

## Abstract

The enteric microbiome and nutrient sensing in the small intestine play critical roles in maintaining host metabolic homeostasis. However, insight is limited to bacteria and fungi. The role of the enteric virome remains poorly understood. Here, we demonstrate that the enteric virome significantly influences carbohydrate digestion and absorption independently of bacteriome. Furthermore, the virome elicits distinct responses across different intestinal cell types. Specifically, it activates programs for carbohydrate digestion and absorption in intestinal epithelial cells, while simultaneously stimulating antigen presenting cells-Th17 cells to produce interleukin-22, a cytokine that curbs excessive carbohydrate uptake. The virome’s effect on carbohydrate digestion and absorption—suppressive or stimulatory—depends on the presence or absence of immune surveillance. This intricate metabolic-immune interplay underscores the enteric virome as a pivotal regulator of host metabolism, and highlights its potential as a therapeutic target for metabolic disorders.

## Introduction

The intestine harbors a complex and diverse microbial community that plays a crucial role in nutrient uptake and immune homeostasis. While extensive research has focused on the bacteriome and mycobiome, recent studies have increasingly highlighted the enteric virome role in maintaining intestinal homeostasis^1,2^. The human virome comprises bacteriophages that infect bacteria, viruses that infect cellular microorganisms such as archaea, and viruses that infect mammalian cells^2^. The intestine is the most abundant site of viral colonization, with viruses outnumbering bacteria by ratios ranging from 1:1 to 10:1^3^. Notably, over 97% of identified enteric viruses are bacteriophages, which influence bacterial growth, metabolism, and virulence, thereby shaping the enteric microbial ecosystem^3^.

Beyond the impact on enteric bacteriome, bacteriophages have been shown to interact with the underlying mammalian cell layers, eliciting cellular metabolic and immune response^4,5^. Enteric bacteriophage composition shifts under various physiological and pathological conditions, particularly in metabolic disorders such as obesity, non-alcoholic fatty liver disease, and type 2 diabetes^1^. Despite advances in sequencing and computational methods for identifying viral genomes in enteric contents, most studies have only reported associations between the virome and diseases, with causal relationships remaining unclear^2^. Moreover, most virome studies rely on relative abundance measurements, often overlooking absolute viral load. Recent insights from bacteriome research suggest that bacterial load, rather than composition alone, plays a critical role in host-bacteriome interactions^6^. In the intestine, bacterial load is closely linked to fecal transit time, pH, and enterotype^6,7^. However, functional understanding of the relationship between enteric viral load and mammalian host interactions are currently lacking. Given that the intestine harbors the largest reservoir of viruses in the body and that nutrient sensing in the small intestine—particularly for carbohydrates and lipids—is closely linked to metabolic health and disorder, a key unanswered question is whether the enteric virome has its own autonomous functionality to contribute to this process^8,9^.

In this study, we investigated the role of the enteric virome in intestinal nutrient sensing across carbohydrate and lipid metabolism. We found that the virome selectively influences carbohydrate metabolism in a bacteria-independent manner, with its effects modulated by antigen presenting cell (APC)-T cell surveillance. Our findings highlight the autonomous role of the virome in metabolic-immune regulation and offer new insights into its complex interplay within the enteric ecosystem.

## Result

### Commensal enteric virome regulates carbohydrate digestion and absorption

To test whether virome participate in nutrient uptake, specific pathogen-free (SPF) mice were fed a chow diet (64% carbohydrate, 12% fat) or a high-fat diet (60% fat, 20% carbohydrate), with or without antiviral cocktail treatment (AVC, Figure 1A, 1B, and Table S1). After 3 days of AVC, the numbers of fecal virus-like particles (VLPs) were significantly reduced, while bacterial colony-forming units (CFUs) remained unchanged (Figure S1A and S1B). Intriguingly, AVC had no effect on body weight, food conversion ratio (FCR), or oral glucose tolerance (OGTT) in 60% HFD-fed mice (Figure 1C and S1C). However, in chow-fed mice, AVC significantly altered these metabolic phenotypes, leading to increased body weight, lower FCR, and impaired OGTT (Figure 1 D and S1D).

**Figure 1.**
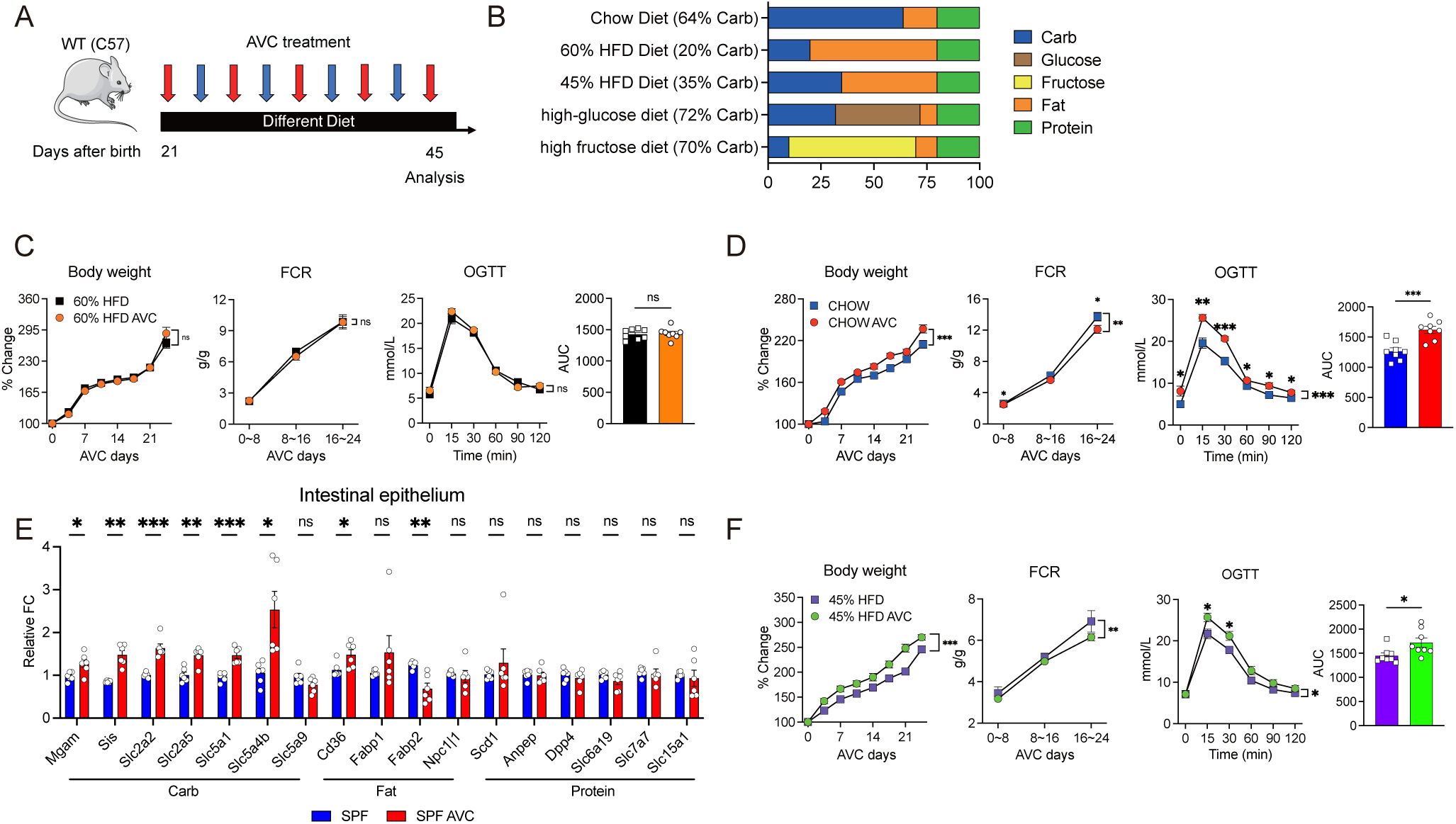
Commensal enteric virome regulates carbohydrate digestion and absorption. (A) Experimental scheme for (C-F). (B) Schematic of the components of each diet, expressed as a percentage of total calories. (C-D) and (F) Body weight, FCR and OGTT in SPF mice on a (C) 60% HFD, (D) chow diet and (F) 45% HFD, treated with or without AVC. n = 4-6 mice/group. (E) qPCR analysis of macronutrient uptake genes transcription in small intestine epithelium of SPF mice on a chow diet, treated with or without AVC. n = 4-6 mice/group. (C-D) and (F) mean ± SEM analyzed by two-way ANOVA with Geisser-Greenhouse correction and two tail t-test. (E) mean ± SEM analyzed by two tail t-test. ∗p < 0.05, ∗∗p < 0.01, ∗∗∗p < 0.001. Data are representative of at least two independent experiments. See also Figure S1.

**Figure S1.**
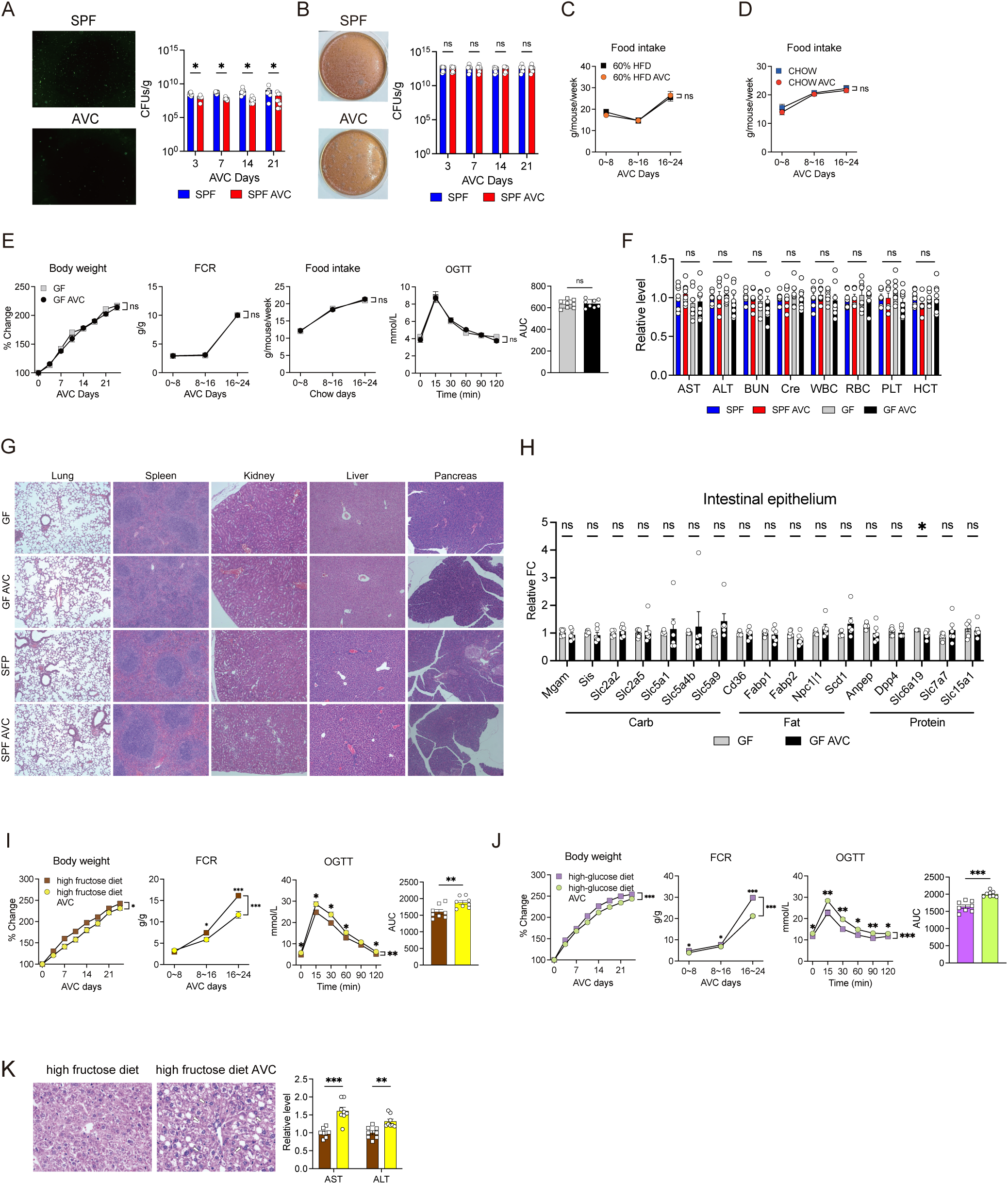
Commensal enteric virome regulates carbohydrate digestion and absorption. Related to Figure 1. (A) Confocal imaging and the counts of SYBR Gold-stained VLPs isolated from SPF mice on a chow diet, treated with or without AVC. n = 4-6 mice/group. (B) Representative images of bacterial culture analysis on CBA plates showing from SPF mice on a chow diet, treated with or without AVC. n = 4-6 mice/group. (C) and (D) Food intake in SPF mice on a (C) 60% HFD and (D) chow, treated with or without AVC. n = 4-6 mice/group. (E) Body weight, FCR, food intake and OGTT in GF mice, treated with or without AVC. n = 4-6 mice/group. (F) AVC does not cause significant changes in AST/ALT and BUN levels, WBC, RBC, or PLT counts, or HCT in both SPF and GF mice. n = 4-6 mice/group. (G) HE staining of organ sections from SPF and GF mice fed a chow diet, treated with or without AVC. n = 4-6 mice/group. (H) qPCR analysis of macronutrient uptake genes transcription in small intestine epithelium in GF mice on a chow diet, treated with or without AVC. n = 4-6 mice/group. (I) and (J) Body weight, FCR and OGTT in SPF mice on a (I) high fructose diet and (J) high-glucose diet, treated with or without AVC. n = 4-6 mice/group. (K) HE staining of liver and AST/ALT in mice fed a high fructose diet, treated with or without AVC. n = 4-6 mice/group. White arrows indicate ballooned hepatocytes. (A) and (B) mean ± SEM analyzed by Mann-Whitney test. (C-E) and (I-J) mean ± SEM analyzed by two-way ANOVA with Geisser-Greenhouse correction and two tail t-test. (F), (H) and (K) mean ± SEM analyzed by two tail t-test. ∗p < 0.05, ∗∗p < 0.01, ∗∗∗p < 0.001. Data are representative of at least two independent experiments.

To exclude the direct drug effects of AVC, we treated germ-free (GF) mice with AVC on a chow diet. Notably, AVC did not alter body weight, FCR, food intake or OGTT in GF mice (Figure S1E). Histological and blood biochemical analyses showed no abnormalities in GF or SPF mice treated with AVC (Figure S1F and S1G). qPCR confirmed that AVC did not alter carbohydrate, lipid or protein digestion and absorption genes transcription in GF mice (Figure S1H). However, in SPF mice, AVC upregulated carbohydrate digestion and absorption (CD&A) genes transcription, while lipid digestion and absorption genes response was non-uniform, and amino acid digestion and absorption remained unchanged (Figure 1E). These findings indicate that AVC-induced metabolic phenotypes are not a direct consequence of drug exposure.

Since AVC selectively affected chow-fed but not 60% HFD-fed mice and mainly influenced CD&A genes, we hypothesized that AVC-induced metabolic phenotypes specifically depend on carbohydrate availability. To test this, we reduced 60% HFD fat content to 45%, compensating with 15% carbohydrates (45% HFD, maltodextrin and sucrose). AVC elicited similar metabolic responses to those observed in chow-fed mice, supporting the carbohydrate-dependence hypothesis (Figure 1F). Since maltodextrin and sucrose are composed of glucose and fructose, we next tested whether AVC-induced metabolic effects depended on carbohydrate type using high-fructose (60% fructose + 10% starch) and high-glucose (32% starch/maltodextrin + 40% glucose) diets. In both cases, AVC lowered body weight but consistently reduced FCR and impaired OGTT (Figure S1I and S1J). Moreover, in high-fructose-fed mice, AVC exacerbated hepatic de novo lipogenesis and elevated aspartate aminotransferase and alanine aminotransferase (AST/ALT) levels, suggesting that AVC aggravated high-fructose diet–induced non-obese steatohepatitis (Figure S1K). As AVC affected metabolic phenotypes in both diets, its impact depended on carbohydrate availability rather than specific carbohydrate type. Together, these results suggest that virome plays a critical role in carbohydrate metabolism.

### Enteric virome regulates CD&A independently of the bacteriome

To determine whether AVC-induced metabolic phenotypes depend on the bacteriome, we disrupted the bacteriome using ampicillin (AMP), either alone or with AVC in chow diet-fed mice (Figure 2A). AMP significantly altered the α- and β-diversity of the bacteriome without affecting CFUs, while AVC robustly reduced VLPs numbers in the AMP context (Figure S2A and S2B). Based on this, mice were divided into four groups: (1) CON (SPF, no AMP, no AVC); (2) AVC (SPF, no AMP, with AVC); (3) AMP (with AMP, no AVC); and (4) AMP AVC (with AMP and AVC). Compared to CON group, AMP group showed no changes in body weight but increased FCR and impaired OGTT, reinforcing the well-recognized role of the bacteriome in metabolic processes^10^ (Figure 2B). However, in the AMP context, AVC still induced metabolic effects similar to those in SPF context, including increased body weight, reduced FCR, and impaired OGTT, suggesting that AVC-induced metabolic phenotypes are likely independent of the bacteriome (Figure 1D and 2B). To further assess the role of the bacteriome, we performed 16S rRNA sequencing and correlation analysis. We first performed a global comparison of overall bacterial community composition across the four groups. Surprisingly, despite AVC impairing metabolism (increased body weight, reduced FCR and impaired OGTT), MicrobiomeAnalyst linked the CON group’s bacterial pattern—rather than the AVC group’s—to metabolic disorders such as fatty liver disease (Figure S2C). Meanwhile, both AMP and AMP AVC groups exhibited bacterial patterns associated with metabolic disorders. These results suggest that overall bacterial community composition alone cannot explain AVC-induced metabolic impairments. We next analyzed individual bacterial correlations with metabolic parameters. In the SPF context, *Intestinimonas*, *Granulicatella*, and *Paramuribaculum* correlated with FCR and OGTT, whereas in the AMP context, *Ruminococcus* and *Clostridia* were associated with these parameters (Figure S2D). Notably, these associations were context-specific, with no bacterial taxa universally linked to AVC-induced metabolic changes. Moreover, the virome may alter bacterial metabolites through horizontal gene transfer without affecting bacterial abundance, while bacterial metabolites influence host metabolism^11^. However, correlation analysis also failed to correlate enteric metabolites with metabolic phenotypes across both contexts (Figure S2E). Furthermore, short-chain fatty acids (SCFAs), key bacterial metabolites in host-bacteriome interactions, remained unchanged following AVC, regardless of the SPF or AMP context (Figure S2F)^12^. To further validate these findings, we tested AVC in the context of imipenem treatment and observed similar results as in the AMP-treated context (Figure S2G). Together, these findings indicate that AVC-induced metabolic phenotypes are independent of the bacteriome. Interestingly, VLPs numbers were strongly correlated with FCR and OGTT (Figure 2C). We previously observed that VLPs numbers declined significantly by day 3 of AVC, while bacteriome diversity remained unchanged (Figure S2H). By this point, OGTT was already impaired, further supporting the potential role of virome load in carbohydrate metabolism (Figure S2I).

**Figure 2.**
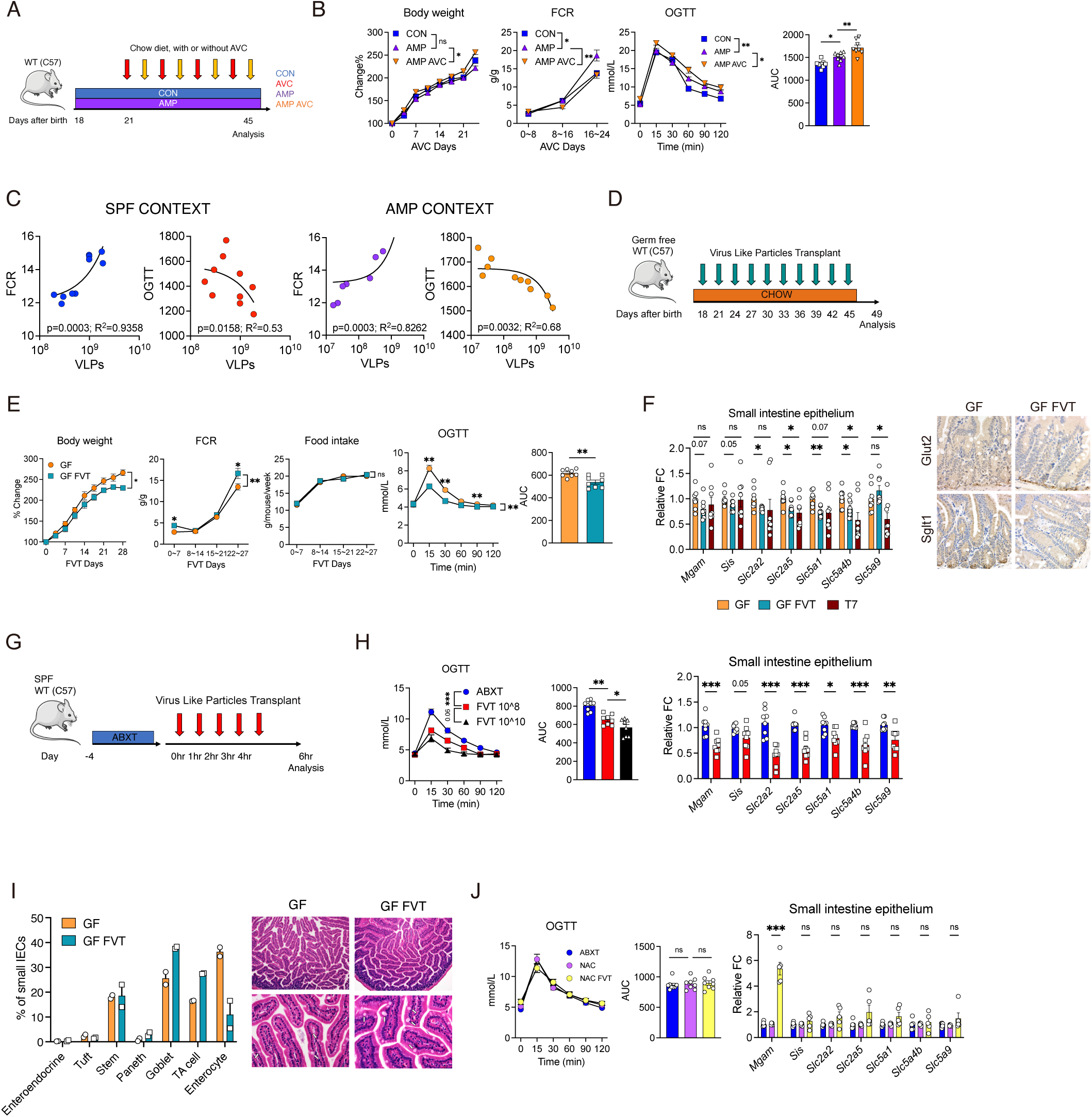
Enteric virome regulates CD&A independently of the bacteriome. (A), (D) and (G) Experimental schemes for (B), (E) and (H) separately. (B) Body weight, FCR and OGTT in AMP mice on a chow diet, treated with or without AVC. n = 4-6 mice/group. (C) Simple linear regression illustrating the Pearson’s correlation between the numbers of VLPs and FCR and OGTT. (E) Body weight, FCR, food intake and OGTT in GF mice on a chow diet, treated with or without FVT. n =4-6 mice/group. (F) Immunohistochemistry staining and qPCR analysis show the CD&A genes expression in small intestine of GF mice treated with or without FVT or T7 transplantation. n = 4-6 mice/group. (H) and (J) OGTT and qPCR analysis of CD&A genes expression in small intestine epithelium (H) of ABXT mice or (J) NAC mice, treated with or without FVT. n = 4-6 mice/group. (I) Frequency of epithelial cell subsets in GF mice treated with or without FVT. White arrows indicate goblet cells. (B), (E), (H) and (J) mean ± SEM analyzed by two-way ANOVA with Geisser-Greenhouse correction and two tail t-test. (F) mean ± SEM analyzed by two tail t-test. ∗p < 0.05, ∗∗p < 0.01, ∗∗∗p < 0.001. Data are representative of at least two independent experiments. See also Figure S2.

**Figure S2.**
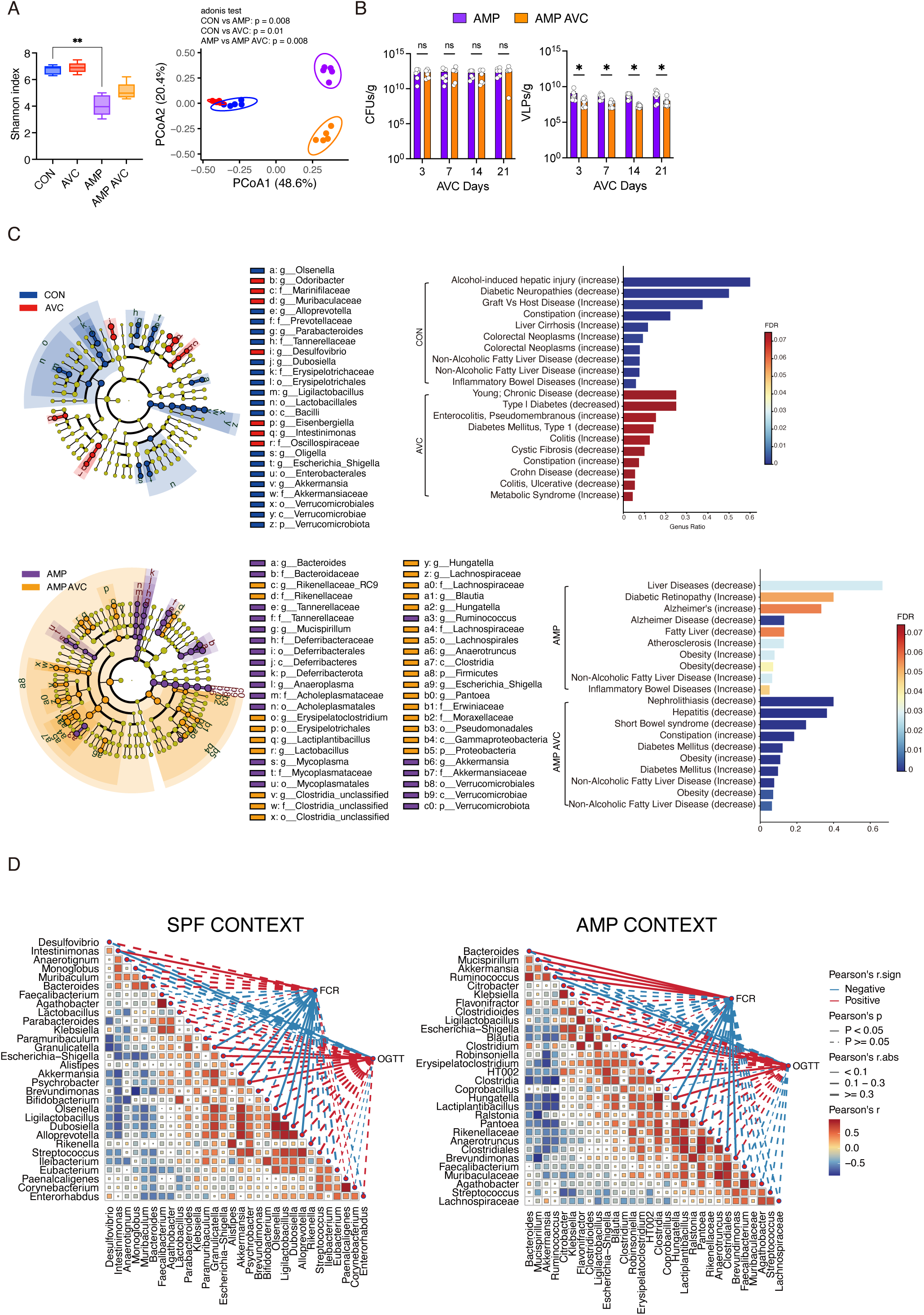
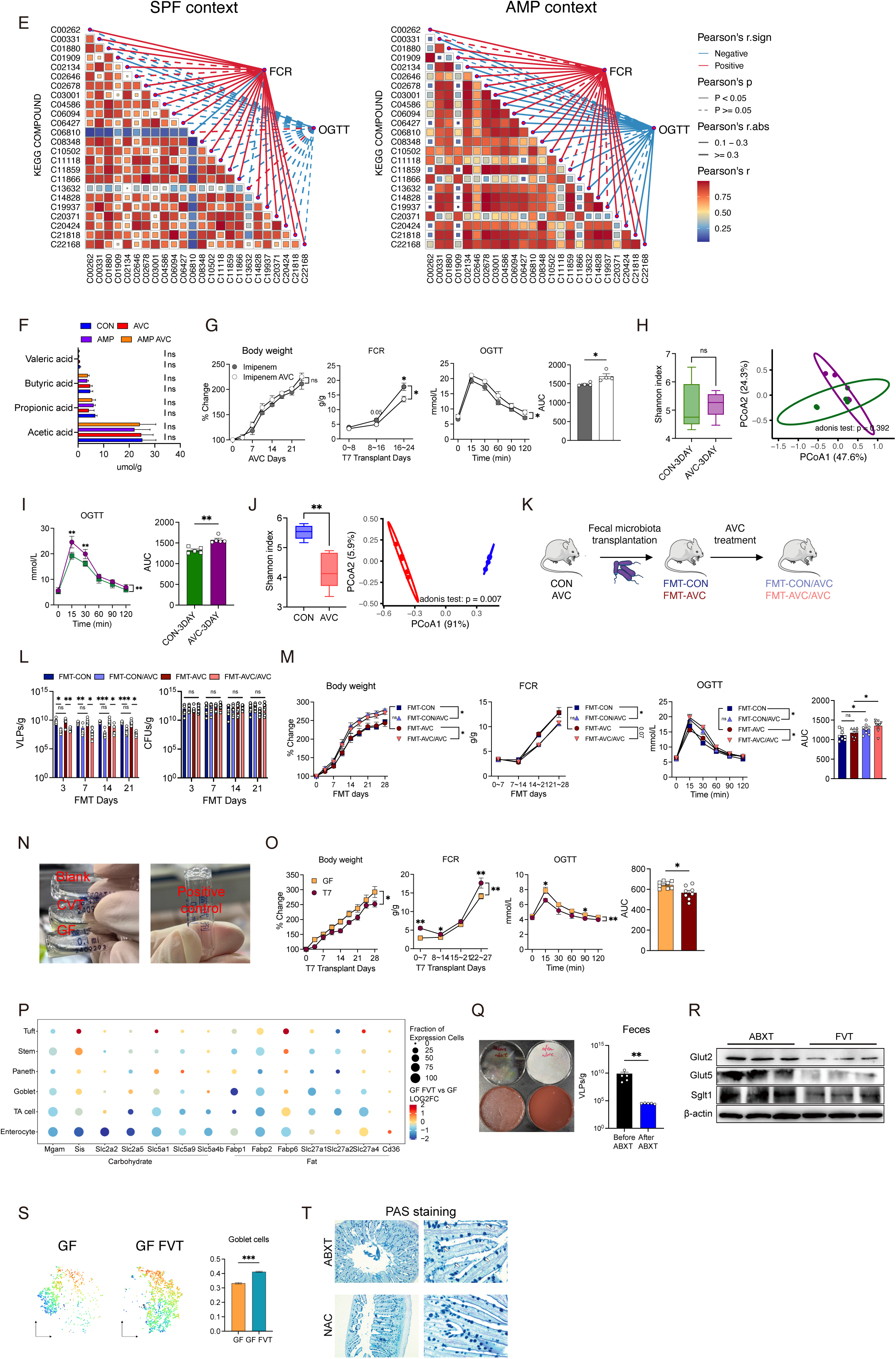
Enteric virome regulates CD&A independently of the bacteriome. Related to Figure 2. (A) Shannon’s diversity and PCoA for enteric bacteriomes from CON and AMP context with or without AVC. Kruskal-Wallis test, followed by Wilcoxon’s rank sum test with Bonferroni’s correction, was conducted for Shannon’s diversity. n = 5 mice/group. (B) and (L) VLPs number and CFUs in the (B) AMP and (L) FMT mice, treated with or without AVC. n = 4-6 mice/group. (C) LEfSe analysis identified the most differentially abundant taxa in the CON and AMP context mice, treated with or without AVC. LDA score > 3 and Wilcoxon< 0.05 are shown. Cladogram of significant changes at all taxonomic levels. The root of the cladogram represents the domain bacteria. The size of node represents the abundance of taxa. Bar chart is used to display the analysis result of MicrobiomeAnalyst. The pool of bacteria was set by LEfSe analysis. (D) and (E) the relationship between (D) bacteria genus or (E) metabolites and FCR and OGTT in CON and AMP context was calculated. The significant levels of Pearson’s correlation among dependent variables were indicated by square size. Network planning showed the Pearson’s correlation between bacteria genus and metabolites and FCR and OGTT. Red lines indicate positive correlations, while blue lines indicate inverse correlations. Edge width corresponds to Pearson’s r value, and edge type indicates statistical significance. Pearson’s correlation coefficient matrix shows the relationships among dependent bacteria genus and metabolites. The significant levels of Pearson’s correlation among dependent bacteria genus and metabolites were indicated by square size. Normality and Lognormality of FCR and OGTT were tested by Kolmogorov-Smirnov test, p>0.1. (F) Levels of SCFA from CON and AMP context mice, treated with or without AVC. n = 5 mice/group. (G) Body weight, FCR and OGTT in imipenem-treat mice on a chow diet, treated with or without AVC. n = 4-6 mice/group. (H) Shannon’s diversity and PCoA for enteric bacteriomes from CON context with or without 3 days AVC. (I) OGTT in mice treated with or without 3 days AVC. n = 4-6 mice/group. (J) Shannon’s diversity and PCoA for enteric viromes from CON context mice, treated with or without AVC. (K) Experimental schemes for (M). (M) Body weight, FCR and OGTT in FMT mice on a chow diet, treated with or without AVC. n = 4-6 mice/group. (N) Representative images of tachypleus amebocyte lysate test for purified VLPs. The inability to form a stable gel indicates that the endotoxin concentration is below the detection limit. (O) Body weight, FCR and OGTT in GF mice on a chow diet, treated with or without T7 transplantation. n = 4-6 mice/group. (P) Bubble plot showing the differential expression of carbohydrate and lipid uptake genes across different epithelial cell subsets. (Q) Representative images of bacterial culture on CBA plates and the numbers of VLPs from fecal samples of mice before and after ABXT. n = 4-5 mice/group. (R) Western blot analysis of CD&A proteins expression in small intestine of ABXT mice, treated with or without FVT. n = 4-6 mice/group. (S) Subset of goblet cells assigned weight of mucus synthesis-related genes (*Muc2, Muc3a, Muc13, Fcgbp, Tff3* and *Clca1*) and results from AUCell analysis. (T) PAS staining of the small intestine of ABXT mice treated with or without NAC. The mucus layer on the surface of intestinal epithelium was significantly reduced after NAC treatment. White arrows indicate mucus. (B), (L) and (Q) mean ± SEM analyzed by Mann-Whitney test. (G), (I), (M) and (O) mean ± SEM analyzed by two-way ANOVA with Geisser-Greenhouse correction and two tail t-test. ∗p < 0.05, ∗∗p < 0.01, ∗∗∗p < 0.001. Data are representative of at least two independent experiments.

Next, we characterized the enteric virome by purifying and sequencing VLPs obtained from enteric content of mice 。Consistent with previous reports, AVC alters both the numbers and composition of the enteric virome (Figure S1A and S2J)^13^. To determine whether virome composition, in addition to its numbers, contributes to AVC-induced metabolic phenotypes, we performed fecal microbiota transplantation (FMT), transferring microbiota from CON and AVC groups into GF mice (FMT-CON and FMT-AVC, Figure S2K). After FMT, without continued AVC, VLPs numbers was restored in both groups (Figure S2L). Under these conditions, no significant differences in body weight, FCR, or OGTT were observed between FMT-CON and FMT-AV groups, suggesting that virome or bacteriome composition is insufficient to drive metabolic phenotypes (Figure S2M). However, reintroducing AVC in FMT mice depleted the VLPs numbers and restored metabolic phenotypes, including increased body weight, reduced FCR, and impaired OGTT (Figure S2K, S2L, and S2M). Together, these findings indicate that AVC-induced carbohydrate metabolic phenotypes are primarily driven by virome load, rather than changes in virome or bacteriome composition.

To investigate whether the virome autonomously drives metabolic phenotype, we enriched VLPs from the enteric contents of SPF mice, ensuring that endotoxin levels in the enriched VLPs were below the detection limit (<0.06 EU/ml, Figure S2N)^14,15^. The enriched 1-5*10^8 VLPs were transplanted into GF mice (Figure 2D). Compared to GF controls, fecal VLPs transplantation (FVT) reduced body weight, increased FCR, and decreased OGTT area under the curve (AUC, Figure 2E). As bacteriophages are the major members of the enteric virome, we transplanted T7 bacteriophages into GF mice, which recapitulated nearly all FVT-induced metabolic effects (Figure S2O). Notably, these observations sharply contrasted with ACV-treated SPF mice, where VLPs depletion led to increased body weight, decreased FCR, and impaired OGTT (Figure 1D). Single-cell RNA sequencing (scRNA-seq) of small intestinal epithelial cells (IECs) revealed suppressed transcription of CD&A genes in the FVT group, consistent with qPCR and immunohistochemistry analyses (Figure 2F and S2P). Given the substantial intestinal physiological differences between GF and SPF mice, we extended these observations to SPF mice treated with antibiotics for gastrointestinal decontamination (ABXT, Figure 2G)^16,17^. ABXT significantly reduced the numbers of VLPs while eliminating intestinal bacteria (Figure S2Q). FVT reduced OGTT AUC and CD&A genes expression in ABXT mice in a dose-dependent manner, mirroring the effect observed in GF mice (Figure 2E, 2F, 2H, and S2R). These findings indicate that the enteric virome autonomously drives metabolic phenotypes independently of the bacteriome.

Beyond CD&A genes transcriptional changes, FVT induced structural remodeling of the intestinal epithelium. VLPs exposure promoted goblet cell hyperplasia and upregulated mucin synthesis-related genes (Figure 2I and S2S). Mucus layers at the surface of intestinal epithelium are primarily composed of mucins, which are predominantly secreted by goblet cells^18^. As the first interface between the host and the external environment, the mucus layer serves as a key site for interactions with the enteric commensal microbiome^18^. Mucins can bind bacteriophages, influencing their transit time in the gastrointestinal tract and potentially mediating their impact on IECs cellular program^19–21^. We next examined whether mucus influences FVT-induced changes in CD&A genes transcription and OGTT. Degrading the mucus layer with N-acetylcysteine (NAC) abolished the suppressive effect of FVT on CD&A genes transcription and OGTT in ABXT mice, underscoring the critical role of mucin-mediated virome interactions in epithelial regulation (Figure 2J and Figure S2T). These findings indicated that mucin serves as a key mediator through which the virome modulates intestinal CD&A.

### Virome regulates CD&A via APCs and T cells

Given that IECs are responsible for CD&A in the small intestine, we investigated whether they can directly sense the virome to suppress CD&A genes transcription. Surprisingly, instead of suppressing CD&A genes transcription, VLPs, T7 bacteriophages, and two mouse-derived purified bacteriophages enhanced it stimulatory program in a dose-dependent manner (Figure 3A and S3A-S3D). This result contrast sharply with our *in vivo* findings, where FVT suppressed CD&A genes transcription in GF and ABXT mice (Figure 2F and 2H). Besides directly stimulating mature enterocyte, FVT may also inhibit CD&A through a mechanism involving the differentiation of specialized enterocytes with a low CD&A expression profile. Transient amplifying (TA) cells, which can rapidly differentiate into mature enterocytes, are crucial for maintaining the absorptive epithelium^22^. However, scRNAseq revealed that VLPs stimulation did not alter CD&A genes transcription in TA cells, suggesting that TA cell differentiation is unlikely to be responsible (Figure S3E).

**Figure 3.**
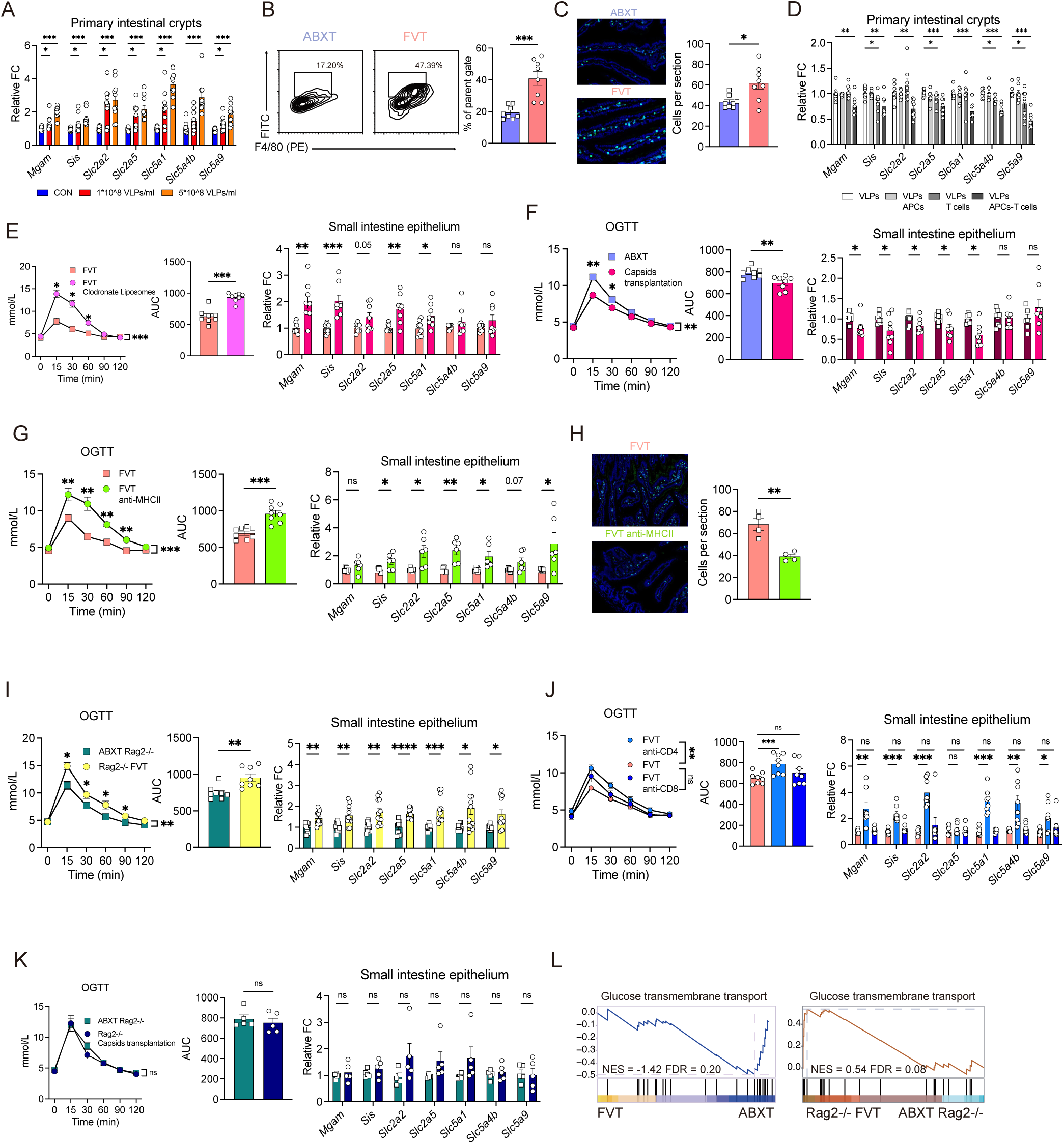
Virome regulates CD&A via APCs and T cells. (A) and (D) represent qPCR analysis of CD&A genes expression in WT intestinal crypts stimulated with (A) VLPs and (D) VLPs with or without APCs or T cells. n = 4-6. (B) Flow cytometry for APCs from the small intestine of fluorescent labeled-VLPs transplant mice. n =4-6 mice/group. (C) and (H) Immunofluorescence staining for CD45 (cyan) and CD3 (green) in the intestine of (C) ABXT mice with or without FVT and (H) FVT mice with or without anti-MHCII. n =4-6 mice/group. (E), (G) and (J) OGTT and qPCR analysis of CD&A genes expression in the small intestines of WT FVT mice treated with (E) clodronate liposomes (G) anti-MHC II antibodies, and (J) anti-CD4 or anti-CD8 antibodies. n = 4–6 mice/group. (F) and (K) OGTT and qPCR analysis of CD&A genes expression in the small intestines of (F) ABXT WT mice and (K) ABXT Rag2-/- mice, treated with or without capsids transplantation. (I) OGTT and qPCR analysis of CD&A genes expression in the small intestines of ABXT Rag2-/- mice treated with or without FVT. n = 4-6 mice/group. (L) GSEA of genes upregulated or downregulated in glucose transmembrane transport in WT-FVT and Rag2-/- FVT. (A-D) and (H) mean ± SEM analyzed by two tail t-test. (E-G) and (I-K) mean ± SEM analyzed by two-way ANOVA with Geisser-Greenhouse correction and two tail t-test. ∗p < 0.05, ∗∗p < 0.01, ∗∗∗p < 0.001. Data are representative of at least two independent experiments. See also Figure S3.

**Figure S3.**
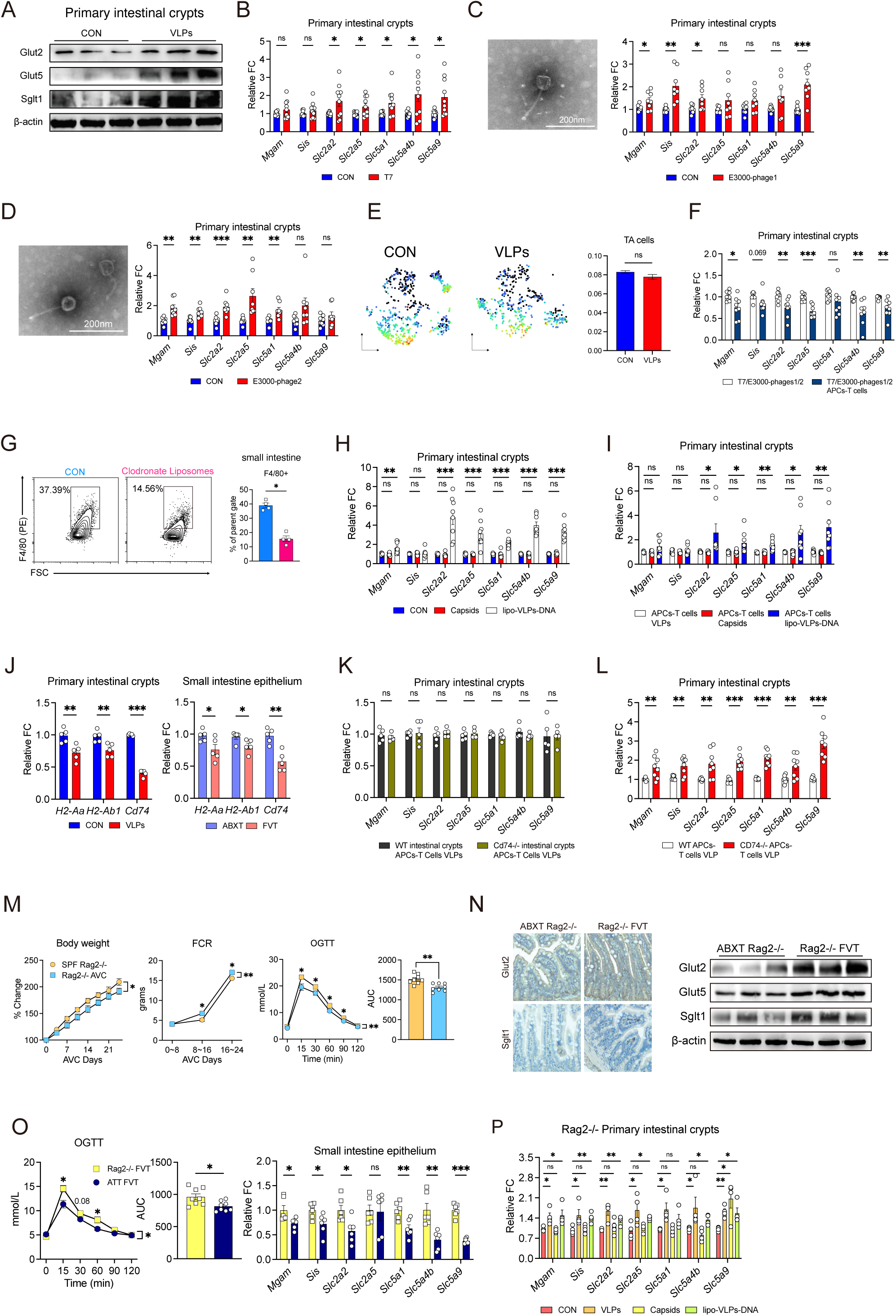
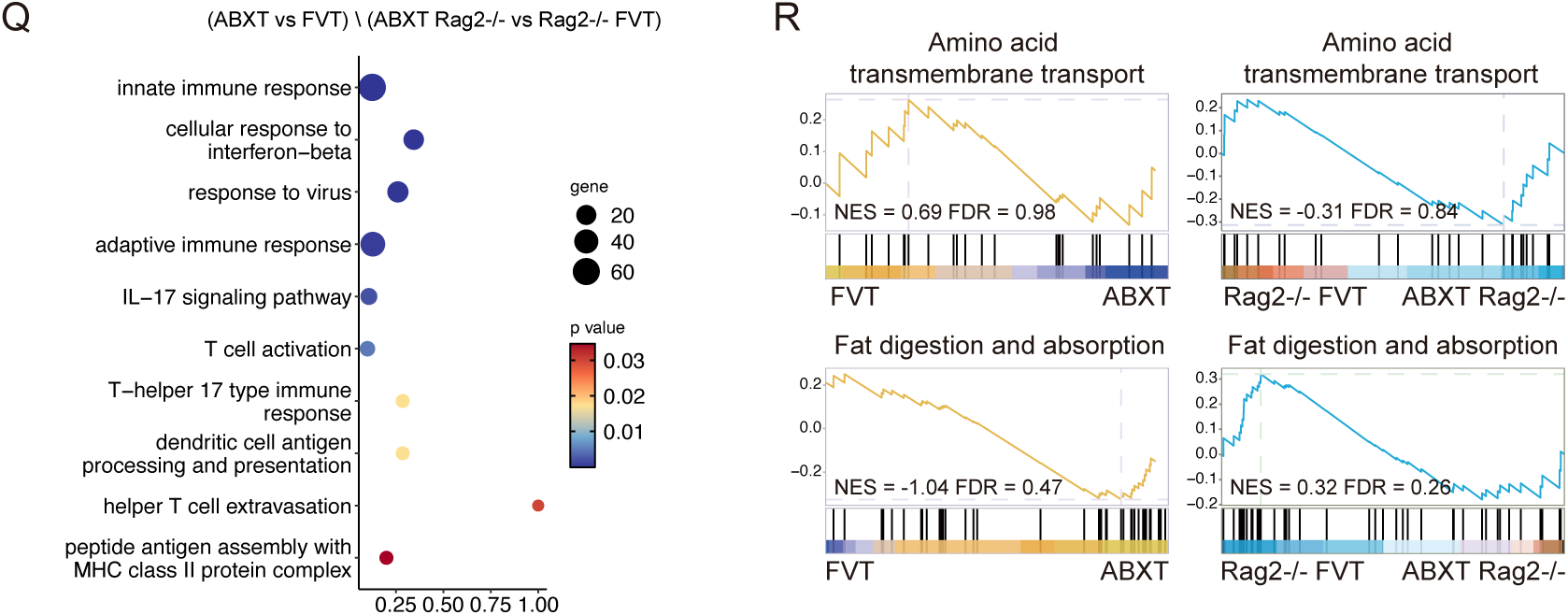
Virome regulates CD&A via APCs and T cells. Related to Figure 3. (A) Western blot analysis of CD&A protein expression in intestinal crypts of WT mice treated with or without VLPs. n = 4-6 mice/group. (B-D) and (H) qPCR analysis of CD&A genes expression in WT intestinal crypts stimulated with or without (B) T7, (C) C3000-phage1, (D) C3000-phage2 (H) capsids or lipo-enveloped VLPs-DNA. n = 4-6. (E) Subset of TA cells assigned weight of CD&A genes and results from AUCell analysis. (F) qPCR analysis of CD&A genes expression in WT intestinal crypts stimulated with T7/E300-bacteriophages, with or without APCs-T cells. n = 4-6. (G) Flow cytometry for CD45+F4/80+ cells from the small intestine of the mice treated with or without clodronate liposomes. n = 4-6 mice/group. (I) qPCR analysis of CD&A genes expression in WT intestinal crypts stimulated with APCs-T cells, with or without capsids or lipo-enveloped VLPs-DNA. n = 4-6. (J) After *in vivo* or *in vitro* VLPs exposure, the expression of genes related to MHC II antigen presentation in IECs was observed. n = 4-6. (K) qPCR analysis of CD&A genes expression in WT or Cd74-/- intestinal crypts stimulated with APCs-T cells-VLPs. n = 4-6. (L) qPCR analysis of CD&A genes expression in WT intestinal crypts stimulated with WT APCs or CD74-/- APC, as well as with T cells and VLPs (n = 4–6). n = 4-6. (M) Body weight, FCR and OGTT in SPF Rag2-/- mice treated with or without AVC. n = 4-6 mice/group. (N) Immunohistochemistry staining of tissue samples and western blot from the small intestine of ABXT Rag2-/- mice treated with or without FVT. (O) OGTT and qPCR analysis of CD&A genes expression in the small intestines of FVT ABXT Rag2-/- mice, treated with or without adoptive T cells transfer. n = 4-6 mice/group. (P) qPCR analysis of CD&A genes expression in Rag2-/- intestinal crypts stimulated with VLPs, capsids or lipo-enveloped VLPs-DNA. n = 4-6. (Q) Bubble chart is used to display the changed genes in GO analysis within the set of (ABXT vs FVT) \ (ABXT Rag2-/- vs Rag2-/- FVT). (R) GSEA of genes upregulated or downregulated in amino acid transmembrane transport and fat digestion and absorption in WT-FVT and Rag2-/- FVT. (B-D), (F-L) and (P) mean ± SEM analyzed by two tail t-test. (M) and (O) mean ± SEM analyzed by two-way ANOVA with Geisser-Greenhouse correction and two tail t-test. ∗p < 0.05, ∗∗p < 0.01, ∗∗∗p < 0.001. Data are representative of at least two independent experiments.

Thus, we hypothesized that other intestinal cells contribute to this suppression. After transplanting fluorescent-labeled VLPs into ABXT mice, we observed that VLPs predominantly accumulated in CD45+F4/80+ cells, coinciding with increased T cell infiltration (Figure 3B and 3C). Immune cells are known to fine-tune intestinal nutrient uptake processes^23^. We next assessed their individual contributions *in vitro*. APCs alone did not inhibit CD&A transcription, while T cells exhibited mild suppression. However, when both were present, they intensified the suppression of all CD&A genes (Figure 3D and Figure S3F). Since VLPs predominantly accumulated in CD45+F4/80+ cells, we next examined APCs *in vivo*. Depleting F4/80+ APCs by clodronate liposomes abolished the suppressive effects of FVT on OGTT and CD&A genes transcription^24^ (Figure 3E and Figure S3G). This confirm that APCs are essential for immune-mediated suppression of CD&A, as T cells alone are insufficient. We next investigated how APCs sense the virome and contribute to immune-mediated suppression. Enteric virome consists primarily of proteins (capsids) and nucleic acids (genome). To systematically examine their contributions, we first tested the effects of these components on IECs. Capsids alone did not alter CD&A genes transcription, whereas lipid-encapsulated nucleic acids mimicked the stimulatory effect of intact VLPs (Figure S3H). Notably, co-culturing APCs and T cells with capsids, but not lipid-encapsulated nucleic acids, reproduced the inhibitory effect on CD&A genes transcription (Figure S3I). *In vivo*, capsid transplantation yielded results similar to FVT (Figure 3F). These findings highlight the critical role of protein capsids in immune-mediated CD&A suppression.

Considering the importance of capsids in APC-T cell regulation of CD&A, we hypothesized that MHC class II (MHCII), which typically presents protein-derived antigens, might mediate APC-T cell communication. Blockade of MHCII *in vivo* abolished the suppressive effects of FVT on CD&A gene transcription and reduced T cell recruitment, indicating the essential role of MHCII in APC-T cell interaction (Figure 3G and 3H). In the small intestine, in addition to APCs, IECs can also express MHCII^25^. Interestingly, VLPs exposure, both *in vivo* and *in vitro*, downregulated MHCII-related antigen presentation genes in IECs (Figure S3J). Moreover, the suppression of CD&A genes transcription by APC-T cell interaction remained unaffected when IECs were derived from Cd74-/- mice (Figure S3K). However, this suppression was diminished when APCs were derived from Cd74-/- mice (Figure S3L). These findings highlight that MHCII expression on APCs, rather than IECs, is key to virome-induced metabolic regulation.

Next, we investigated the role of T cells *in vivo*. In Rag2-/- mice, AVC induced a distinct metabolic phenotype compared to WT mice. Instead of increased body weight, lower FCR and impaired OGTT, Rag2-/- mice treated with AVC exhibited lower body weight, higher FCR, and reduced OGTT AUC (Figure S3M). Whereas FVT suppressed CD&A genes transcription and improved OGTT in ABXT WT mice, it instead led to upregulated CD&A genes transcription and impaired OGTT in ABXT Rag2-/- mice (Figure 3I and S3N). Importantly, adoptive transfer of T cells into Rag2-/- mice restored the inhibitory effects of FVT on OGTT and CD&A genes transcription in the small intestine epithelium (Figure S3O). To pinpoint the specific subset of T cells involved, we selectively depleted CD4+ or CD8+ T cells using blocking antibodies. CD8+ T cell depletion did not alter CD&A genes transcriptions, whereas CD4+ T cell depletion abolished the inhibitory effects of FVT (Figure 3J). These results highlight the crucial role of T cells, particularly CD4+ T cells, in mediating the virome-induced metabolic regulation.

Notably, in the Rag2-/- mice, VLPs directly stimulated CD&A genes transcription, consistent with our *in vitro* findings (Figure 3A and 3I). Direct stimulation of IECs from both WT and Rag2-/- mice with intact VLPs increased CD&A genes transcription, an effect dependent on nucleic acids but not on capsid (Figure S3H and S3P). Consistently, capsids transplantation failed to reproduce the stimulatory effect of FVT in Rag2-/- mice, suggesting nucleic acids drive this effect (Figure 3K). While we cannot confirm whether nucleic acids promote CD&A upregulation in Rag2-/- mice by directly stimulating IECs, these findings nevertheless highlight the functional heterogeneity of viral components in regulating CD&A.

To further explore the virome’s effect on small intestinal genes expression and immune-dependent effects, we performed bulk RNA sequencing on WT and Rag2-/- mice, with or without FVT. Sequencing data identified key differentially expressed genes linked to immunity, antigen presentation, antiviral responses, and helper T cell function pathway (Figure S3Q). GSEA analysis revealed a genotype-dependent effect of FVT on CD&A. FVT suppressing CD&A in WT mice but enhancing it in Rag2-/- mice (Figure 3L). Instead, FVT did not alter protein or lipid metabolism in either genotype (Figure S3R). Together, these results highlight the virome’s role in regulating CD&A in the small intestine, with its effect—suppressive or stimulatory—dependent on the presence of absence of APC-T cell surveillance.

### Virome suppresses CD&A through Th17 cell-IL-22

Considering the pivotal role of CD4+ T cells in virome-mediated suppression of CD&A, we assessed whether FVT modulates T cell-associated cytokine receptor expression in the small intestine. In WT mice, FVT upregulated the interleukin-22 (IL-22) receptor, whereas such changes were not observed in Rag2-/- mice (Figure S4A). GSEA analysis revealed that IL-22 receptor-associated pathways, including STAT protein phosphorylation and PI3K-Akt signaling, were activated exclusively in FVT-treated WT mice (Figure S4B). ELISA and qPCR confirmed that IL-22 and *Il22ra1* expression were elevated in the intestines of FVT-treated WT mice but remained unchanged in Rag2-/- mice (Figure 4A and S4C). In the intestine, IL-22 is primarily produced by group 3 innate lymphoid cells (ILC3) and Th17 cells^23^. FVT selectively enhanced IL-22 production from Th17 cells but not from ILC3 in WT mice (Figure 4B and Figure S4D). Supporting this, FVT did not alter *Il22* or *Il22ra1* expression in Rag2-/- or CD4+ T cell-depleted mice, which retain ILC3 but lack Th17 cells (Figure S4C and S4E). Microbial stimuli have been reported to drive ILC3 toward ex-ILC3, characterized by reduced IL-22 secretion and increased T-bet and IFN-γ expression^26,27^. Similarly, VLPs stimulation upregulated T-bet and IFN-γ in ILCs (Figure S4F and S4G). IL-22 is a well-established regulator of intestinal epithelial metabolism. Consistently, IECs treated with IL-22 showed dose-dependent downregulation of the CD&A genes transcription, confirming that this program is negatively regulated by IL-22 (Figure 4C). *In vivo*, IL-22 blockades reversed the FVT-induced suppression of OGTT and upregulated CD&A genes transcription (Figure 4D). Conversely, IL-22 supplementation in Rag2-/- mice reduced the AUC of OGTT and downregulated CD&A genes transcription (Figure 4E). Together, these findings establish IL-22 as a key mediator through which the virome regulates small intestinal CD&A via immune cells.

**Figure 4.**
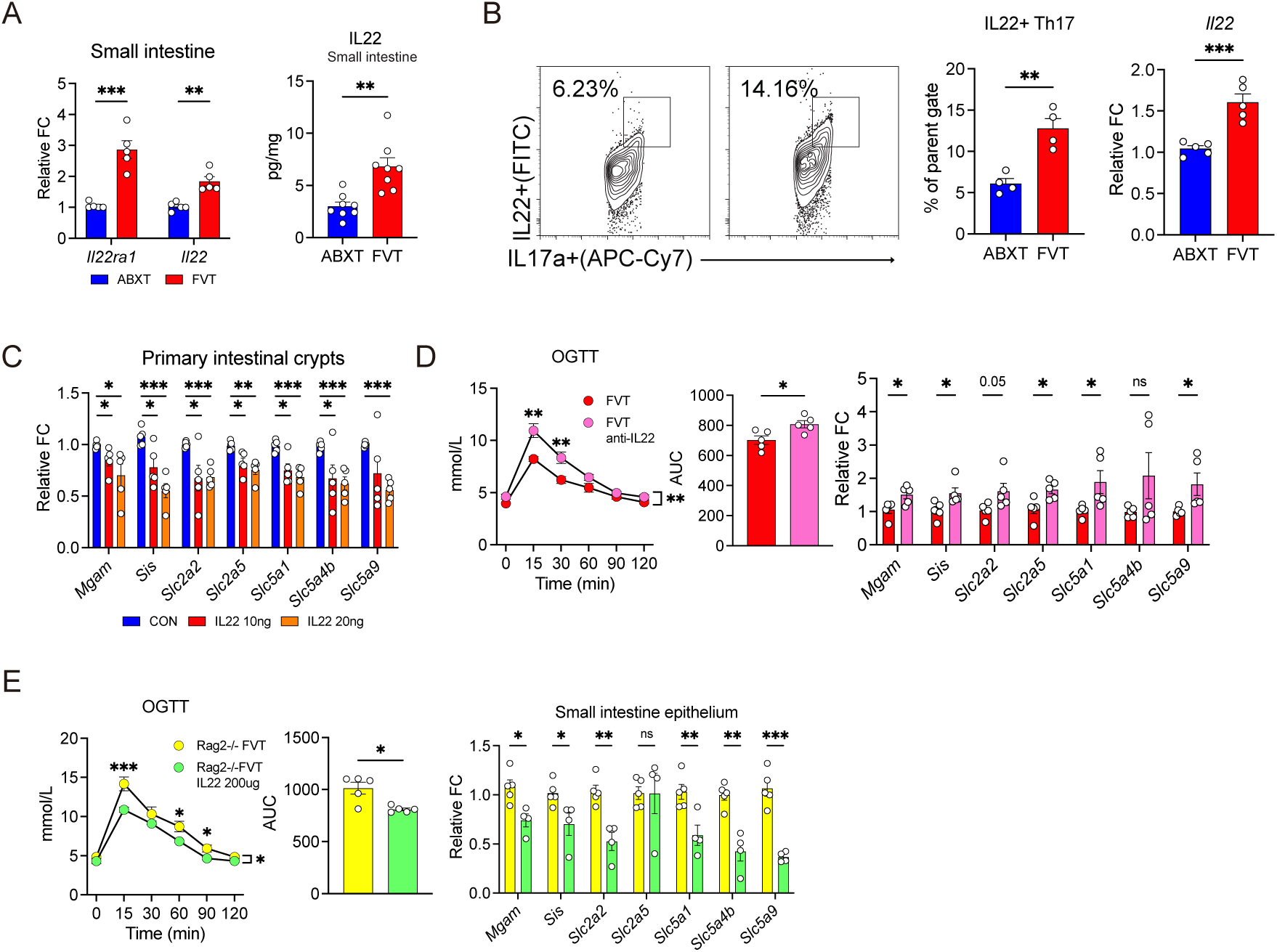
Virome suppresses CD&A through Th17 cell-IL-22. (A) ELISA and qPCR analysis of IL-22 and *Il22ra1* in the small intestine of WT mice treated with or without FVT. n = 4-6 mice/group. (B) Flow cytometry of IL-22 expression in Th17 from WT mice treated with or without FVT. n = 4-5 mice/group. (C) CD&A genes expression in intestinal crypts stimulated with or without IL-22 recombinant protein. n = 4-5. (D) OGTT and qPCR analysis of CD&A genes expression in small intestine in WT FVT mice treated with or without IL-22 antibody. n = 4–6 mice/group. (E) OGTT and qPCR analysis of CD&A genes expression in small intestine in Rag2-/- FVT mice treated with or without IL-22 recombinant protein. n = 4-6 mice/group. (A-C) mean ± SEM analyzed by two tail t-test. (D) and (E) mean ± SEM analyzed by two-way ANOVA with Geisser-Greenhouse correction and two tail t-test. ∗p < 0.05, ∗∗p < 0.01, ∗∗∗p < 0.001. Data are representative of at least two independent experiments. See also Figure S4.

**Figure S4.**
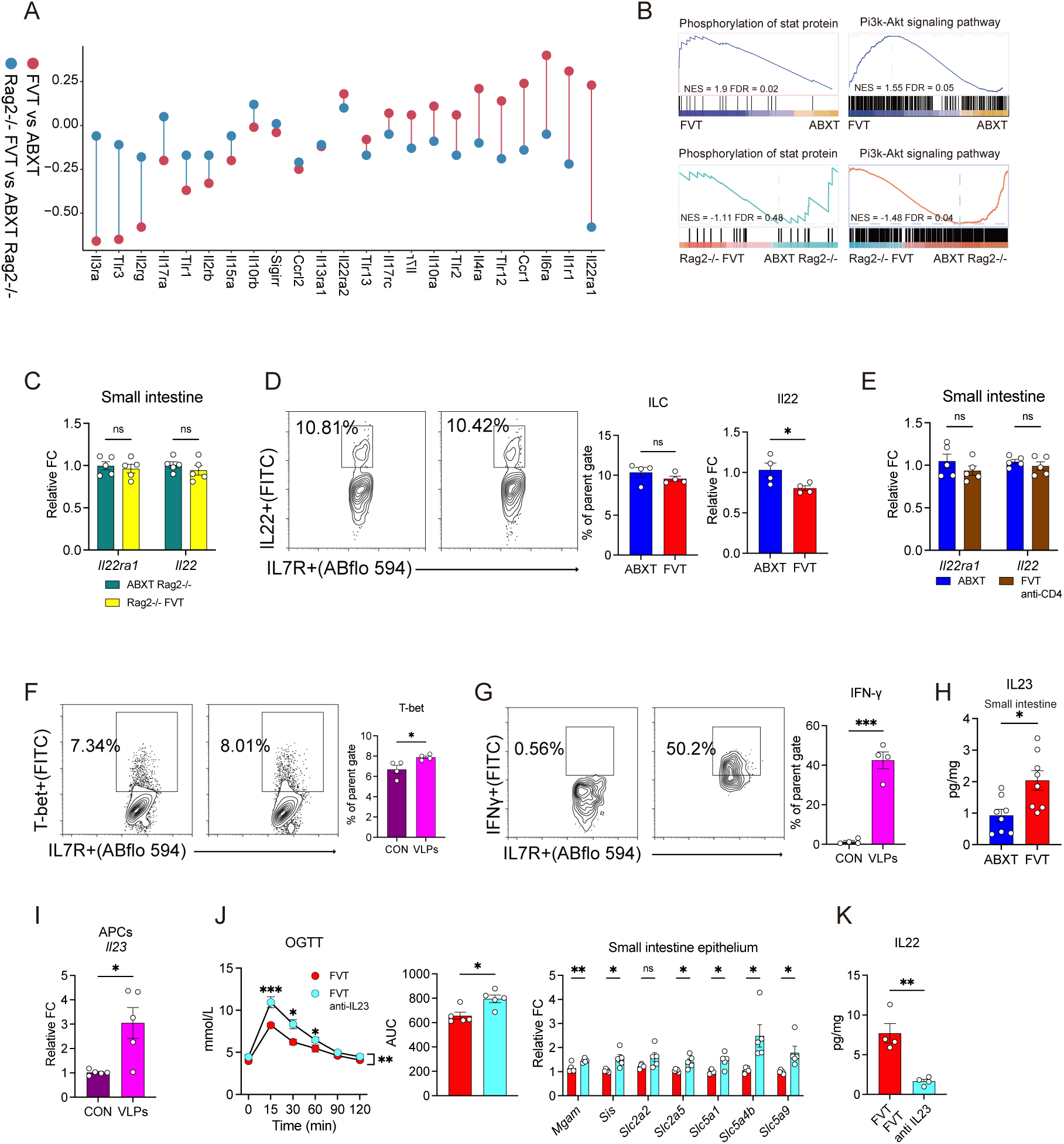
Virome suppresses CD&A through Th17 cell-IL-22. Related to Figure 4. (A) Dumbbell plot is used to display the changed T cells-associated cytokine receptor genes transcription in WT FVT and Rag2-/- FVT mice. (B) represents GSEA of genes upregulated or downregulated in IL-22 related signal pathway in WT and Rag2-/- mice, treated with or without FVT. (C) and (E) qPCR analysis of *Il22* and *Il22ra* in the small intestine of (C) Rag2-/- mice and (E) CD4+ T cells depletion mice, treated with or without FVT. n = 4-6 mice/group. (D) Flow cytometry of IL-22 expression in ILCs from WT mice, treated with or without FVT. n = 4-5 mice/group. (F) and (G) Flow cytometry of (F) T-bet and (G) IFNγ expression in ILCs stimulated with VLPs. n = 4-5. (H) ELISA analysis of IL-23 in the small intestine of WT mice treated with or without FVT. n = 4-5 mice/group. (I) qPCR analysis of *Il23* transcription in APCs treated with or without VLP. n = 4-5/group. (J) OGTT and qPCR analysis of CD&A genes expression in small intestine in WT FVT mice, treated with or without IL-23 antibody. n = 4–5 mice/group. (K) ELISA analysis of IL-22 in the small intestine of WT FVT mice, treated with or without IL-23 antibody. n = 4-5 mice/group. (C-I) and (K) mean ± SEM analyzed by two tail t-test. (J) mean ± SEM analyzed by two-way ANOVA with Geisser-Greenhouse correction and two tail t-test. ∗p < 0.05, ∗∗p < 0.01, ∗∗∗p < 0.001. Data are representative of at least two independent experiments.

We previously established the critical role of APCs in suppressing CD&A, and IL-22 secretion by Th17 cells is primarily dependent on IL-23 produced by APCs^28,29^. ELISA confirmed that FVT-treated mice exhibited increased IL-23 protein levels (Figure S4H). *In vitro*, exposure to VLPs also increases IL-23 expression in APCs (Figure S4I). IL-23 blockades reversed the FVT-induced suppression of OGTT and restored CD&A genes transcription, accompanied by decreased IL-22 expression (Figure S4J and S4K).

## Discussion

Our study has discovered an underappreciated virome-host communication at the interface of the metabolic and immune system. We found that the enteric virome influences the CD&A in a bacteria-independent manner. The virome autonomously promotes CD&A program in intestinal epithelial cells, while simultaneously stimulating APCs and Th17 cells to produce IL-22, which counteracts excessive CD&A. The virome’s effect on CD&A—suppressive or stimulatory—depends on the presence or absence of APC-T cell surveillance.

The enteric microbiome is a known regulator of nutrient metabolism, but the functional role of the virome remains unclear, as most studies rely on metagenomic associations rather than functional evidence^9,14^. Distinguishing the autonomous role of the virome is challenging due to its close interplay with the bacteriome. To address this, we utilized FVT in GF and ABXT models to separate the virome from bacterial influences and directly assess its impact. There are multiple protocols for isolating VLPs, each with their own biases and limitations. Given the abundance of bioactive metabolites in enteric contents that regulate host metabolism, we first precipitated VLPs with PEG, then used chloroform to remove remain bacterial cells and excess PEG^14,15^. We deemed this step essential to avoid contaminating metabolites that would confound subsequent experiments.

Our study reveals a clear dose-dependent role of the gut virome in regulating host immunity and metabolism. This highlights the importance of considering not only compositional differences but also viral load—a critical yet underappreciated dimension—in future functional studies of the virome. While absolute quantification strategies have begun to emerge in bacterial microbiome research, virome studies still largely rely on relative abundance measures^6,7,30,31^. Notably, viral abundance in the gut is not static but shaped by diverse physiological and pathological conditions. For example, patients with Crohn’s disease exhibit a marked expansion of *Caudovirales* phages, and in infants, VLPs undergo a rapid postnatal increase followed by a gradual decline around two years of age, indicating age-dependent fluctuations^14,32–37^. Our findings further demonstrate that antiviral agents such as AVC and acriflavine effectively reduce gut VLP abundance, supporting the notion that viral load is a regulated and functionally relevant biological variable. We propose that viral quantity itself constitutes a meaningful biological signal, with its impact on glycemic control contingent upon immune surveillance. Incorporating absolute virome abundance into analytical frameworks will be essential for decoding virome-mammalian interactions and for integrating viral ecology into metabolic and immune models.

Given the dominance of bacteriophages in enteric virome, interplay between the bacteriome and virome is expected^38,39^. However, the autonomous role of the virome in regulating host metabolism is unclear. Much evidence demonstrates that bacteriophages are readily internalized by mammalian cells and directly modulate cellular processes independent of their bacterial host^4,5,21,36,40–42^. Research on bacteriophages recognition by mammalian cells has primarily focused on nucleic acid-sensing pathways such as RIG-I and cGAS^14,36,43^. However, some studies suggest that cellular changes can also occur even without detectable activation of these pathways, indicating the involvement of other viral components^4^. Our findings provide new data on this issue, demonstrating that virome recognition is cell type-specific and viral component-dependent. Specifically, viral nucleic acids mediate the stimulatory CD&A program of the virome on epithelial cells, while viral capsid proteins are the key drivers of immune-mediated suppressive regulation. Importantly, MHCII antigen presentation is essential for virome-induced immune suppression of CD&A. These findings not only underscore the importance of considering both viral components and cell types in virome-host interactions, but also suggest that bacteriophage capsids may function as microbial protein antigens. Indeed, bacteriophage-specific antibodies are frequently observed in circulation, and bacteriophage capsids can be processed via MHC pathways in APCs^42,44–47^. Given the abundance and diversity of the enteric virome, further elucidating how the innate and adaptive immune systems recognize these commensal viruses will provide valuable insights into immune regulation.

Prior studies have established that ILC3-Th17-IL-22 axis play a pivotal role in intestinal homeostasis, with bacterial signals regulating this balance^48–53^. Our findings provide new insights, showing that, like bacterial stimuli, virome stimulation can induce ILC3 transition to an ex-ILC3 state, leading to the loss of IL-22 secretion^26,27,54,55^. However, virome also activates APC-T cell interactions, leading to increased IL-22 production. This compensatory mechanism counteracts the loss of ILC3-derived IL-22, maintaining glucose homeostasis by limiting excessive CD&A. These results highlight the enteric virome as an active participant in immune regulation.

In conclusion, our study establishes the enteric virome as an active regulator of host metabolism, functioning beyond its conventional role in shaping bacteriome. By integrating metabolic and immune signaling, the virome autonomously influences nutrient uptake and immune homeostasis. Future therapeutic and nutritional strategies for metabolic disorders should incorporate the enteric virome as a significant factor of intervention.

## METHODS

### Animal

All animal experiments conducted in this study were approved by the Experimental Animal EthICs Committee of Jinling Hospital affiliated to Medical School of Nanjing University. C57BL/6J and Rag2-/- (Strain NO. T011534) mice were sourced from Nanjing GemPharmatech Co. Cd74-/- (NM-KO-190772) mice were sourced from Shanghai Model Organisms Center, Inc. The mice were housed in SPF facilities, maintained in a temperature-controlled environment at 26°C with a 12-hour light/dark cycle to minimize external variability. Sterile C57/BL6J mice were bred and housed in a sterile mouse facility at Nanjing GemPharmatech Co. GF status monitoring: VLPs and mice feces were inoculated into three liquid culture media—FTM, BHI, and TSB. The FTM and BHI cultures were incubated at 37°C and transferred onto CBA plates at 7 and 14 days, followed by microscopic examination. CBA agar plates were incubated at 37°C for 48 hours to check for bacterial growth. TSB cultures were incubated at 25°C for 14 days to detect fungal growth. The GF status of the samples was determined based on the growth results across the three-culture media and microscopic examination. Mice aged 3 weeks were utilized for the AVC and GF-FVT experiments, while those aged 6-8 weeks were used for other experiments. All feeds were irradiated and sterilized. Mice had *ad libitum* access to food and water. The AVC and AMP treatment was following the method as previously described^9,43^. In brief, mice were treated with antiviral cocktail containing ribavirin (30mg/kg), lamivudine (10mg/kg) and acyclovir (20mg/kg), or AMP (1g/L) in drinking water. For ABXT, mice received sterilized water supplemented with AMP (10 mg/mL), neomycin trisulfate salt hydrate (10 mg/mL), metronidazole (10 mg/mL), and vancomycin hydrochloride (5 mg/mL), with a volume of 10 μL/g body weight^56^. NAC (30 mM, 200 μL per mouse) was orally administered for four days^57^. The concentration of imipenem added to the drinking water was 0.25 g/L. Detailed dietary compositions are provided in Table S1. All animal experiments included both male and female mice equally, with cells and tissues used for bulk RNA sequencing and sc-RNA seq derived from male mice.

### Enteric content and feces bacterial culture

Fresh fecal samples or intestinal contents were collected and weighed. The samples were then diluted 1:10,000 in sterile water. A 100 μL aliquot of the diluted sample was plated onto CBA plates and incubated anaerobically at 37°C for 48 hours. CFUs were subsequently counted.

### Fecal microbiota transplantation

Fresh fecal samples and intestinal contents were collected and immediately transferred to an anaerobic chamber for processing. One gram of mixed stool was resuspended in 5 mL of PBS containing 10% glycerol and thoroughly homogenized. The suspension was then filtered through a 70 μm cell strainer and stored at −80 °C until use. For FMT administration, 150 μL of the fecal suspension was delivered to GF mice via oral gavage.

### Glucose tolerance test

After a 12-hour fast, mice underwent an OGTT, where glucose was administered orally at a dosage of 2g/kg. Blood samples were obtained from the tail vein at 0, 15, 30, 60, 90, and 120 minutes post-glucose challenge for glucose measurement using Accu-Chek glucose meters^58^.

### Enumeration of VLPs

The enteric contents were resuspended in SM buffer at a ratio of 1:5 (SM buffer: 8 mM magnesium sulfate heptahydrate, 50 mM Tris-Cl, 100 mM NaCl, and 0.01% gelatin)^14^. After homogenization for 5 minutes, the mixture was centrifuged at 2700 g for 15 minutes. The supernatant was collected and centrifuged at 25000 g for 15 minutes, repeating until no precipitate was observed. Next, 0.2 volume of peg buffer (peg buffer: 200g/L peg8000, 2.5M NaCl) was added, and the mixture was left in an ice bath overnight. The following day, centrifugation was performed at 3300 g for 1 hour. The supernatant was discarded, and the precipitate was resuspended in PBS buffer. Then, 0.3 volume of chloroform was added, gently mixed, and centrifuged at 2500g for 5 minutes and collect the aqueous phase. The aqueous phase was then incubated with DNase (DN-25, Sigma) 1mg/ml at 37°C for 1 h to remove DNA unprotected by capsids. And heat inactivated DNase at 65°C for 15 minutes. A portion of the VLPs was plated onto CBA plates and incubated anaerobically for 24 hours to check for bacterial contamination. Endotoxin levels were assessed according to the manufacturer’s instructions (BK-T04, BOKANG BIO).

### Histological examination

The organs samples were fixed in 4% phosphate-buffered formaldehyde, paraffin-embedded, and subsequently stained with hematoxylin and eosin.

### Immunofluorescent and Immunohistochemistry Assay

For Immunofluorescent staining, slides were incubated overnight at 4°C in a humid chamber with antibodies against CD3 and CD45 (1:200 dilution, ABclonal), followed by incubation with biotinylated secondary antibody (1:500 dilution, ABclonal) for one hour. Images were captured using Olympus FV3000 confocal microscope and Olympus OlyVIA system^59^. For Immunohistochemical staining, the small intestine tissues were fixed in 10% neutral-buffered formalin and embedded in paraffin; 5-µm sections were affixed to slides, and then deparaffinized and stained with Glut2 and Sglt1 antibodies (20436-1-AP and 30861-1-AP, Proteintech).

### Quantification of VLPs

Adapted from the method by Mária Džunková et al^60^, bacteriophage T7 and bacteriophage MS2 were stained with SYBR Gold (S11494, Thermo Fisher) at 4°C for 30 minutes as a positive control. Ultrapure water was filtered through a 100 kDa ultrafiltration tube (UFC5100, Millipore), and the filtrate was used as a negative control. The FITC channel sensitivity of the flow cytometer was set to the highest, and the cell size was set to the lowest. T7, MS2, and filtrate were detected at the slowest flow rate, and bacteriophage gates were drawn. VLPs were extracted using the aforementioned method. The VLPs were then stained with SYBR Gold for 30 minutes and dialyzed through a 100 kDa ultrafiltration membrane overnight. The dialyzed VLPs were detected by fluorescence microscope and flow cytometer. Particles with sizes <0.5 µm were considered VLPs.

### Intestinal crypt isolation and treatment

The mouse small intestines were longitudinally cut and cleaned with PBS. Subsequently, the small intestine was cut into 5mm pieces and placed in pre-cooled 10mM EDTA/PBS buffer at 4°C, incubated on ice for 30 minutes. After vigorous shaking for 5 minutes, the mixture was allowed to stand for 5 minutes, and the supernatant was discarded. The precipitate was resuspended in 10mM EDTA/PBS buffer and subjected to another 30-minute ice bath^61^. After shaking again for 5 minutes, 5% FBS was added, followed by a 5-minute incubation period. The supernatant was filtered through a 70μm cell sieve (Millipore). The purity and density of intestinal crypts were observed under a microscope. If the content of intestinal crypts was low, the precipitation was resuspended with 10mM EDTA/PBS buffer and the steps were repeated until clean intestinal crypts were obtained. The intestinal crypts were then centrifuged at 100g for 2 minutes, resuspended in FBS, and used for subsequent experiments^61^.

### Isolation of intestinal lymphocyte, T cells and APCs

The mouse small intestines were longitudinally cut and cleaned with PBS. Subsequently, the intestine was cut into 5 mm pieces and placed in pre-cooled 5 mM EDTA/PBS buffer at 4°C for 30 minutes on ice. After vigorous shaking for 5 minutes, the suspension was centrifuged at 300 g for 5 minutes, and the supernatant was discarded^62^. The precipitate was then resuspended in collagenase III (100 U/ml, LS004182, Worthington) and incubated in a water bath at 37°C for 10 minutes. To stop the action of collagenase III, 5% FBS was added. The resulting cell suspension was filtered through a 40 μm cell sieve (Millipore) and centrifuged at 300 g for 5 minutes. After resuspending the pellet in 35% Percoll (P7828, Sigma Aldrich), it was layered slowly onto 75% Percoll for density centrifugation. Centrifugation at 700 g for 30 minutes resulted in the collection of the milky white cell layer^63^, from which T cells or F4/80+APCs were further extracted using a isolation kit as per experimental requirements (130-095-130 and 130-110-443, Miltenyi Biotec).

### Coculture of APCs, T cells and Intestinal crypts

After co-incubating APCs with VLPs overnight, the cells were washed twice with PBS. An equal number of T cells were then added and co-incubated. Subsequently, 5*10^5^ T cells-APCs and intestinal crypts, derived from whole intestine extracts of one mouse, were co-incubated for 2 hours at 37°C and 5% CO_2_ in a total volume of 1ml FBS.

### Cytokine Detection

Intestinal tissues were collected from euthanized mice, homogenized and centrifuged, and the supernatant was stored at −20 °C for ELISA analysis of IL-22 and IL-23. A Mouse IL-22 High Sensitivity ELISA Kit (EK222HS, Multi sciences) and Mouse IL-23 ELISA Kit (EK223, Multi sciences) were utilized to measure the protein concentrations of IL-22 and IL-23 proteins level according to the manufacturer’s instructions.

### T cell, F4/80+ APC depletion, cytokine depletion and IL-22 recombinant protein treatment *in vivo*

T cells were depleted using 250 µg of anti-CD4, anti-CD8 antibodies (BE0003-41, BE0004-1, BioXcell), while F4/80+ APCs were eliminated with clodronate liposomes (40337ES08, Yeasen) following the manufacturer’s instructions. For cytokine neutralization, 100 µg of anti-IL-22 (16-7222-82, Thermo Fisher) or anti-IL-23 (p19) (BE0313, BioXcell) was administered. To block MHCII antigen presentation, 500 µg of anti-MHC Class II (I-A/I-E) (BE0108, BioXcell) was injected. Corresponding isotype controls were used for each treatment group. Mice were intraperitoneally injected with 100 μg of Mouse IL-22 recombinant protein (210-22, Thermo) daily for two consecutive days before the FVT experiment.

### Adoptive T cell transfers

For adoptive transfer experiments, 1 million T cells purified from the spleen and mesenteric lymph nodes were intravenously transferred into Rag2-/- mice. ABXT and FVT treatment was performed one week later.

### Single-Cell Suspension Preparation

Isolated intestinal crypts were filtered through a 70 μm cell strainer and then gently agitated in TrypLE Express (12604039, Thermo Fisher) for 1 minute. The resulting cell suspension was filtered through a 40 µm cell strainer, centrifuged at 150 g for 3 minutes, and finally resuspended in 5% FBS-PBS.

### Single cell RNA sequencing

Single-cell suspensions were processed using the 10x Chromium system to capture approximately 5,000 individual cells, following the manufacturer’s protocol for the 10x Genomics Chromium Single-Cell 3’ Kit (v3). Subsequent cDNA amplification and library construction were carried out according to the standard procedure. The libraries were sequenced on an Illumina NovaSeq 6000 platform (paired-end, 150 bp reads) with a minimum sequencing depth of 20,000 reads per cell.

### qPCR analysis

Tissue or cell was processed with the Trizol reagent (15596018CN, Thermo Fisher) according to the manufacturer’s instructions to extract total RNA^64^. Subsequently, the RNA was reverse transcribed to cDNA using a kit (AG11728, Accurate biology). Relative quantification of gene expression was performed using the comparative CT method, with *Rpl13* serving as the internal controls (RR820Q, Takara). Primer sequences are provided in Table S8.

### Extraction of VLPs protein capsids

After obtaining VLPs as described above, an 0.2 volume of 0.5 M EDTA (ST069, Beyotime) was added and incubated at 65°C for 1 h. Cool on ice and then add DNase (DN-25, Sigma) 1mg/ml at 37°C for 1 h. And heat inactivated DNase at 95°C for 15 minutes^14^.

### Extraction of VLPs DNA for lipo-enveloped experiment

After obtaining VLPs as described above, an equal volume of 0.5 M EDTA was added. DNA extraction was then performed using Phenol/Chloroform/Isoamyl alcohol (25:24:1) (P3803, Sigma Aldrich) according to the manufacturer’s instructions. Following precipitation with ammonium acetate at -20°C overnight, the supernatant was removed by centrifugation at 16,000g for 30 minutes. The resulting pellet was resuspended in 70% ethanol, centrifuged at 16,000g for 3 minutes, and the supernatant was discarded. The pellet was then dried at 65°C for 3 minutes, and the DNA was resuspended in TE buffer^65^. VLP-DNA was encapsulated using lipofectamine 2000 (Thermo Fisher) according to the manufacturer’s instructions.

### Viral DNA sequencing

The previously described method was employed to enrich VLPs, followed by purification of E.Z.N.A.® DNA Kit (D3892-01, Omega Bio-tek), in accordance with the–manufacturer’s instructions. Subsequently, library construction was conducted using the total DNA and TruSeq Nano DNA sample preparation Kit from Illumina, as per the manufacturer’s protocol (15041877, Illumina). Libraries were then size selected for DNA target fragments of ∼400bp on 2% Low Range Ultra Agarose followed by PCR amplified using Phusion DNA polymerase (M0530S, New England Biolabs) for 15 PCR cycles. Next, all samples were sequenced by the Illumina NovaSeq 6000 platform. The raw paired end reads were trimmed and quality controlled by Trimmomatic. Host sequences were removed using the BWA tool, and the Megahit tool was utilized to assemble the remaining sequences. For the assembled genome, sequences > 2000bp were selected, and those sequences were performed virus identification using deepVirFinder (VirFinder score > 0.7, p value < 0.05), VirSorter2 (default), VIBRANT (default), IMG/VR (identify >= 90% & cove rage >= 75%), geNomad (default) and phabox (default) database. vOTUs were clustered using Mummer software to compare candidate virus sequences (similarity > 95%, coverage > 85%, the longest representative one). The novelty and completeness were check by blastn (max target seqs > 5, e value < 1e-5, outfmt < 6, num threads > 8) and checkv (default)^66–68^.

### Concentration and fluorescent labeling of VLPs

After extracting the VLPs, the dialysis sacks with MWCO of 100kDa were used to dialyze the VLPs suspension at 4℃ overnight, transfer the dialysate to Amicon® Ultra centrifuge ultrafiltration tube (UFC5100, Millipore) for concentration. Measure the protein concentration of the concentrate, and control the protein concentration at 2-20mg/mL. Use DMSO to dissolve 5(6)-Carboxyfluorescein N-hydroxysuccinimide ester (C107884, aladdin) to 10mg/mL, slowly add 50-100µL 5(6)-Carboxyfluorescein N-hydroxysuccinimide ester reagent to the 1ml protein sample, mix thoroughly, and incubate at room temperature for 60min while stirring. The labeled VLPs suspension was used for subsequent experiments.

### Bacteriophage plaque assays

Mouse feces were homogenized in 1 mL of PBS and filtered through a 0.45 µm filter to obtain the filtrate. The filtrate was then incubated with C3000 for 3 days, followed by plaque assay for phage detection and subsequent phage purification^69^.

### Bacterial DNA extraction and sequencing

Enteric contents underwent DNA extraction using the CTAB method as per the manufacturer’s instructions^70^. The extracted total DNA was eluted in the elution buffer and subjected to PCR analysis. Subsequently, the samples were sequenced on an Illumina NovaSeq platform according to the manufacturer’s instructions. Paired-end reads were sorted based on their unique barcode and then trimmed by removing the barcode and primer sequence. FLASH was utilized to merge the paired-end reads. Quality filtering of the raw reads was performed under specific conditions to obtain high-quality clean tags using fqtrim. Chimeric sequences were eliminated using Vsearch software. After dereplication using DADA2, a ASV feature table and feature sequence were obtained. Feature sequences were annotated with SILVA database.

### RNA sequencing and analysis

Total RNA extraction was performed using Trizol reagent following the manufacturer’s protocol. Transcriptome libraries were constructed using the TruSeqTM RNA Sample Preparation Kit from Illumina and sequenced on the Illumina NovaSeqTM 6000 platform, generating 2 x 150 bp paired-end reads^71^. Clean reads were aligned to the reference genome by Hisat2 and Genome^72^. Differential expression genes were identified by calculating the expression level of each transcript using the fragments per kilobase of exon per million mapped reads method^73^.

### Western blot analysis

Frozen intestinal crypts were homogenized in RIPA lysis buffer (Beyotime, P0013B) supplemented. After centrifugation of the lysates, protein concentration was estimated in the supernatant using a BCA Protein Assay Kit (23250, Thermo Fisher). Total proteins (30 μg) were separated on 10% SDS-polyacrylamide gel and then transferred to Immobilon®-P PVDF membrane (Sigma Aldrich, IPVH00010). Membranes were blocked with 5% milk in TBS 1× Tween 0.1% for one hour before being probed overnight at 4°C with primary antibodies. After washing, membranes were incubated with secondary antibodies. Signals were recorded by the Tanon 5200.

### Statistics

Statistical analyses, excluding sequencing analysis, were done using GraphPad Prism 10, with details provided in figure legends.

